# Lateral hypothalamus CRFR1 regulation of chronic binge drinking: divergence along anterior-posterior axis

**DOI:** 10.1101/2025.09.16.676507

**Authors:** Jobe Ritchie, Maya Eberle, Jessica Wojick, Madison Campeau, Lili Kooyman, Todd Thiele, Thomas Kash

**Affiliations:** Bowles Center for Alcohol Studies, University of North Carolina at Chapel Hill, Chapel Hill, NC, USA; Department of Psychology and Neuroscience, University of North Carolina at Chapel Hill, Chapel Hill, NC, USA; Department of Pharmacology, University of North Carolina at Chapel Hill, Chapel Hill, NC, USA

**Keywords:** Binge alcohol drinking, corticotropin-releasing factor, lateral hypothalamus

## Abstract

Binge alcohol drinking increases the risk of developing an alcohol use disorder (AUD) and comorbid psychopathology. The lateral hypothalamus (LH) is a brain structure that integrates cognitive and sensory information to tightly regulate motivated behavior, including binge drinking. Importantly, LH function is vulnerable to modulation by the pro-stress neuropeptide corticotropin-releasing factor (CRF), and acute antagonism of CRF receptor 1 (CRFR1) in the LH blunts binge drinking. However, the role of LH CRFR1 in chronic binge drinking is unknown. We used genetically targeted knockdown (KD) of CRFR1 in the LH of male and female mice followed by three weeks of binge drinking using the “Drinking in the Dark” (DID) model. CRFR1 KD in the posterior LH increased alcohol consumption, independent of sex, with no effect of KD in the anterior LH. Consistent with this, total alcohol consumption was negatively correlated with the location of CRFR1 KD in the LH along the anterior-posterior axis. CRFR1 KD did not alter water consumption or body weight, suggesting the effects of CRFR1 KD on alcohol consumption were not due to broad disruption of fluid intake or homeostatic function. In contrast to the observed effects on binge drinking, CRFR1 KD increased anxiety-like behavior and blunted sucrose preference, independent of KD location in the LH. Our findings provide foundational insight into LH function in the context of AUD and prompt further investigation into the divergent roles that distinct circuitry or cell populations along the anterior-posterior axis of the LH may play in binge drinking.

**Highlights:**

- CRFR1 knockdown in posterior LH increased alcohol drinking independent of sex
- Alcohol intake negatively correlated with AP location of CRFR1 knockdown in LH
- CRFR1 knockdown was anxiogenic with more pronounced effects in females
- Sucrose preference was blunted by CRFR1 knockdown

## INTRODUCTION

Excessive alcohol intake is a major public health concern that causes a significant social and economic burden. According to the 2023 National Survey on Drug Use and Health, 21.7% of people aged 12 or older in the United States reported binge drinking during the past month^1^, with binge drinking defined as a pattern of alcohol consumption leading to a blood alcohol concentration ≥ 0.08% (80 mg/dL)^2^. Approximately one third of all alcohol related deaths are associated with binge drinking^3^, and the economic cost of binge drinking accounts for an estimated 76.4% of the total cost of alcohol misuse^4^. Binge drinking also puts individuals at greater risk for the development of an alcohol use disorder (AUD) and associated long term health and productivity consequences^5^. Thus, understanding the mechanisms that support the development or escalation of binge drinking is critically important from a risk prevention and treatment perspective.

Cycles of binge drinking and withdrawal lead to engagement of pro-stress neurotransmitter systems, including corticotropin-releasing factor (CRF)^6^. Binge alcohol consumption acutely increases CRF immunoreactivity in stress-associated structures such as the extended amygdala and paraventricular nucleus of the hypothalamus^7^, and increased CRF signaling through CRF receptor 1 (CRFR1) is a major factor contributing to binge alcohol consumption in rodent models. Disruption of CRFR1 signaling by centrally administered antagonists or brain-wide genetic deletion blunts binge alcohol consumption^8,9^. Binge drinking is particularly reliant on CRFR1 signaling, as CRFR1 antagonism in the central amygdala (CeA) decreases binge-drinking, without altering non-binge consumption of alcohol^7^. CeA CRFR1 antagonism also robustly blunts the escalated alcohol consumption observed during alcohol withdrawal^10^. Notably, the CeA has dense output to other pro-stress brain regions critical for regulating binge alcohol intake, including the lateral hypothalamus (LH).

The LH integrates reward and stress signaling to orchestrate appetitive behaviors. A well-established literature has characterized the strong control the LH exhibits over consummatory behaviors, with lesion of the LH blunting eating and drinking^11^. Similarly, electrical stimulation of the LH robustly drives feeding behavior. LH function is regulated by pro-stress peptidergic signaling through multiple CRF inputs from stress-associated brain regions^13,14^. We recently reported that acute pharmacological blockade of CRFR1 in the LH, as well as chemogenetic silencing of the CRF-positive CeA to LH neuronal circuit, blunts binge-like alcohol consumption in males, but not females^15^. These findings implicate LH CRFR1 in regulation of an acute bout of binge drinking, but it is unclear if this role extends to chronic binge drinking.

There is neuroanatomical and functional divergence of the LH along the anterior-posterior extent. Despite early evidence of the distinct neurocircuitry of the anterior versus posterior LH (aLH, pLH, respectively)^16^, only recently have investigators begun characterizing the function of the LH with respect to its underlying anatomical diversity^17^. The aLH and pLH have differing glutamatergic projections, with the aLH primarily containing projections to the lateral habenula (LHb), while the pLH primarily innervates the ventral tegmental area (VTA)^17^. Notably, the expression of CRFR1 also differs across the anterior-posterior extent of the LH, with more CRFR1 mRNA found in the pLH^18^. Based on the neuroanatomical differences between aLH and pLH, CRFR1 signaling in these structures may play distinct roles in regulation of motivated behaviors. However, the role of aLH/pLH CRFR1 in chronic binge drinking remains to be assessed.

The present study investigated the role of LH CRFR1 expression in binge alcohol consumption, anxiety-like behaviors, and sucrose preference. To that end, male and female CRFR1^fl/fl^ mice received GFP-control or Cre-virus infusions directed at the LH to knock down (KD) CRFR1 expression. Mice were then assayed for anxiety-like behaviors, binge alcohol consumption using the Drinking in the Dark (DID) model, and sucrose preference. Variability in viral targeting along the anterior-posterior axis and recent evidence of functional divergence in LH function and neuroanatomy along this axis prompted analysis of the data based on anterior-posterior (AP) distance from bregma. This led to the identification of divergence in the effects of CRFR1 KD, including effects that are specific to alcohol consumption. The following data provide strong evidence for differences in CRFR1 function in the aLH and pLH in the context of binge drinking and prompt future investigations into the unique underlying neurocircuitry of the LH.

## METHODS

### Animals

Adult male (n = 33) and female (n = 30) CRFR1^fl/fl^ mice^15^ (age 8-12 weeks at start of experiment) were bred in-house and group-housed with littermates prior to the start of the experiment and were then singly housed in ventilated cages in a colony room on a 12:12 h reverse light-dark cycle (lights off at 7 AM). Mice were allowed *ad libitum* access to food (5P76 Irradiated RMH 3000 Rodent Diet) and water unless otherwise stated. All procedures were approved by the Institutional Animal Care and Use Committee of the University of North Carolina at Chapel Hill and performed in accordance with the National Institute of Health’s Guide for the Care and Use of Laboratory Animals.

### Surgery

Mice were anesthetized with 4% isoflurane and placed in a stereotaxic frame (Kopf Instruments). Bilateral craniotomies were performed above the injection site. Hamilton syringes were used to inject 200 nl of AAV8-hSyn-EGFP (GFP control; Addgene; 4.4 x 10^12^) or AAV8-hSyn-Cre-EGFP (Cre; Addgene; 4.5 x 10^12^) at a rate of 100 nl/minute bilaterally into the LH (AP: −1.0, ML: ±1.1, DV: −5.1). Injectors were left in place for 5 minutes to allow for diffusion. During recovery, mice received daily injections of meloxicam (5 mg/kg) for 3 d or until preoperative body weight was reached. Experiments began four weeks post-surgery to allow for virus expression.

### Open Field Test

The open field test (OFT) was conducted in a 51 cm x 51 cm opaque white plexiglass box with an open top. The open field was illuminated from above by an LED strip (center lux of 100). Mice were placed in the corner with their heads facing the center. The session was recorded on a Logitech web cam, and time spent in the zones of the maze were calculated using Noldus behavior tracking software. A 70% ethanol solution was used to clean the open field between subjects.

### Elevated Plus Maze

The elevated plus maze (EPM) assay was conducted on a plus shaped plexiglass maze measuring 76 cm x 76 cm consisting of two walled arms (closed) and two unwalled arms (open). The maze was illuminated from above by two dim LED strips (open arm lux of 25) mounted to the ceiling. Mice were placed in the center of the apparatus with their heads facing the opening of the closed arms and allowed to explore for 10 minutes. The session was recorded on a Logitech web cam and time spent in the zones of the maze were calculated using Noldus behavior tracking software. A 70% ethanol solution was used to clean the maze between subjects.

### Drinking in the Dark

Binge drinking in mice was modeled using a two-bottle choice drinking in the dark (DID) procedure. Mice were allowed access to a 20% (v/v) EtOH solution bottle and a water bottle four days per week, for three weeks. Sessions began three hours into the dark phase of the light/dark cycle and were two hours long on Monday-Wednesday, and four hours long on Thursday. Bottles were weighed before and after the sessions. Average drip values were calculated within cohort and subtracted from daily drinking volumes. Water and ethanol consumption were normalized to body weight (ml/kg) and used to calculate a preference score, dividing the total volume of EtOH consumed (g/kg) by the total volume of fluid consumed (EtOH + water). After the final four-hour drinking session, ∼10 µL of trunk blood was collected from each animal. These blood samples were centrifuged at 4°C for 10 minutes at 2000 RCF, and blood alcohol concentrations (BAC) were calculated using the Analox Alcohol Analyzer (AM1, Analox Instruments).

### Home Cage Sucrose Preference

Mice were given access to water and 1% sucrose in water solution for four hours, at least 7 days after the final DID session. Bottles were weighed at the 2- and 4-hour time points. Water and sucrose consumption were normalized to body weight (ml/kg) and used to calculate a preference score, dividing the total volume of sucrose consumed (g/kg) by the total volume of fluid consumed (sucrose + water).

### Perfusion and Histology

Mice were anesthetized using tribromoethanol (Avertin, 250 mg/kg, intraperitoneal) and transcardially perfused with 25 mL phosphate buffered saline (PBS) followed by 25 mL 4% paraformaldehyde (PFA) solution. Brains were post fixed for 24 hours in 4% PFA solution and then stored in PBS. Brains were sliced into 40 µM sections on a vibratome, mounted onto SuperFrost plus slides, and coverslipped using mounting medium containing DAPI. Virus placement in the LH along the anterior/posterior (AP) plane was assessed on an Echo confocal microscope at 10x magnification (San Diego, CA). The slice containing the brightest labeling was used to define viral placement and each mouse was assigned an AP bregma value based on the closest approximate section/location in the Franklin & Paxinos mouse brain atlas^19^. Mice were then grouped into aLH and pLH using AP −1.35 as the center (≤−1.34 = pLH). Placement was assessed by observers blind to treatment conditions.

### Fluorescence in situ hybridization

Mice were anesthetized using isoflurane and rapidly decapitated. Brains were quickly removed and transferred to dry ice. Frozen brains were covered in Optimal Cutting Temperature (OCT) Compound (Fisher Scientific) and stored at −80°C until they were sliced on a cryostat to produce 16 μm coronal sections containing the LH. Sections were adhered to Superfrost Plus microscope slides and immediately refrozen before being stored at −80°C. Following the manufacturer’s protocol for fresh frozen tissue for the V2 RNAsope manual assay (Advanced Cell Diagnostics), slides were fixed for 15 min in ice-cold 4% PFA solution and then dehydrated in a sequence of ethanol serial dilutions (50, 70, and 100%) and stored overnight in 100% ethanol in −20°C. The following day, slides were briefly air dried and then a hydrophobic barrier was drawn around the tissue sections using a Pap pen (Vector Labs). Slides were then incubated with hydrogen peroxide solution for 10 min, washed in distilled water, and treated with the protease IV solution for 30 min at room temperature. Following protease treatment, C2 and C3 cDNA probe mixtures specific for mouse tissue were prepared at a dilution of 50 diluent:1:1 probe using the following probes from Advanced Cell Diagnostics: GFP (C3, 409011-C3), *Crhr1* (C2, 418011-C2). Sections were incubated with cDNA probes (2 hours) in a humidifying chamber at 40°C, and then underwent a series of signal amplification steps using FL v2 Amp1 (30 min), FL v2 Amp 2 (30 min), and FL v2 Amp3 (15 min) at 40°C in a humidifying chamber in the HybEZ oven (ACD Bio). Two min of washing in 1x RNAscope wash buffer was performed between each step. Sections then underwent fluorophore staining via treatment with a series of TSA plus horseradish peroxidase (HRP) solutions, and Opal 520 and 690 florescent dyes (1:3000 dilution, PN FP1487001KT, PN FP1497001KT). All HRP solutions were applied for 15 minutes and Opal dyes for 30 minutes at 40°C in a humidifying chamber, with an additional HRP blocker solution added between each iteration of this process for 15 min at 40°C and rinsing of sections between all steps with the wash buffer. Last, sections were stained for DAPI using the reagent provided by the Fluorescent Multiplex kit and then coverslipped with Prolong Diamond Antifade mounting media and left to dry overnight at 4°C. Sections from mice were used with a probe for bacterial mRNA (dapB, ACD Bio, 310043) to serve as a negative control that displayed no fluorescent staining. Slices containing the LH were imaged at 10x magnification on an Echo confocal microscope. Cell detection and positive cell quantification were performed using QuPath software^20^.

### Data Analysis

Mice with unilateral or misplaced virus expression were excluded from analysis (n = 11). Statistical analyses were performed using SPSS statistics software. Data were analyzed using Analysis of Variance (ANOVA) or mixed factorial ANOVA, with sex (male, female), virus (GFP, Cre), and AP bregma (aLH, pLH) as between-subjects factors and time/day as within-subjects factors, or with t-tests, where appropriate. Significant interaction effects were followed up using Bonferroni’s *post hoc* test. Significance was set at *p* = 0.05.

## RESULTS

### Genetically targeted CRFR1 knockdown in the LH

The role of LH CRFR1 expression in anxiety-like behaviors, binge alcohol consumption, and sucrose preference was assessed using viral mediated knockdown of CRFR1 (**Fig. 1A**). Male and female CRFR1^fl/fl^ mice received bilateral intra-LH injections of a GFP control virus (n = 17 male, 14 female) or GFP-Cre virus (n = 16 male, 16 female) to knockdown LH CRFR1 (**Fig. 1B-C**). Histological validation of virus expression was conducted to identify the site of injection, and mice were assigned an AP value relative to bregma based on the nearest approximate section in the Franklin & Paxinos mouse brain atlas^19^ (**Fig. 1D**). All groups were approximately equally represented along the anterior-posterior axis. Mice were then grouped into anterior or posterior LH (aLH, pLH; Bregma ≤ −1.34 assigned as posterior) for subsequent analysis. Cre-virus expression resulted in a ∼29% reduction of CRFR1 mRNA in GFP^+^ cells in the LH **(Fig. S1A-E**; t-test, GFP vs Cre, percent GFP^+^CRFR1^+^, *t* = 3.08, *p* = 0.04), in line with published literature using the same constructs^15^, confirming the validity of our knockdown approach.

**Figure 1.**
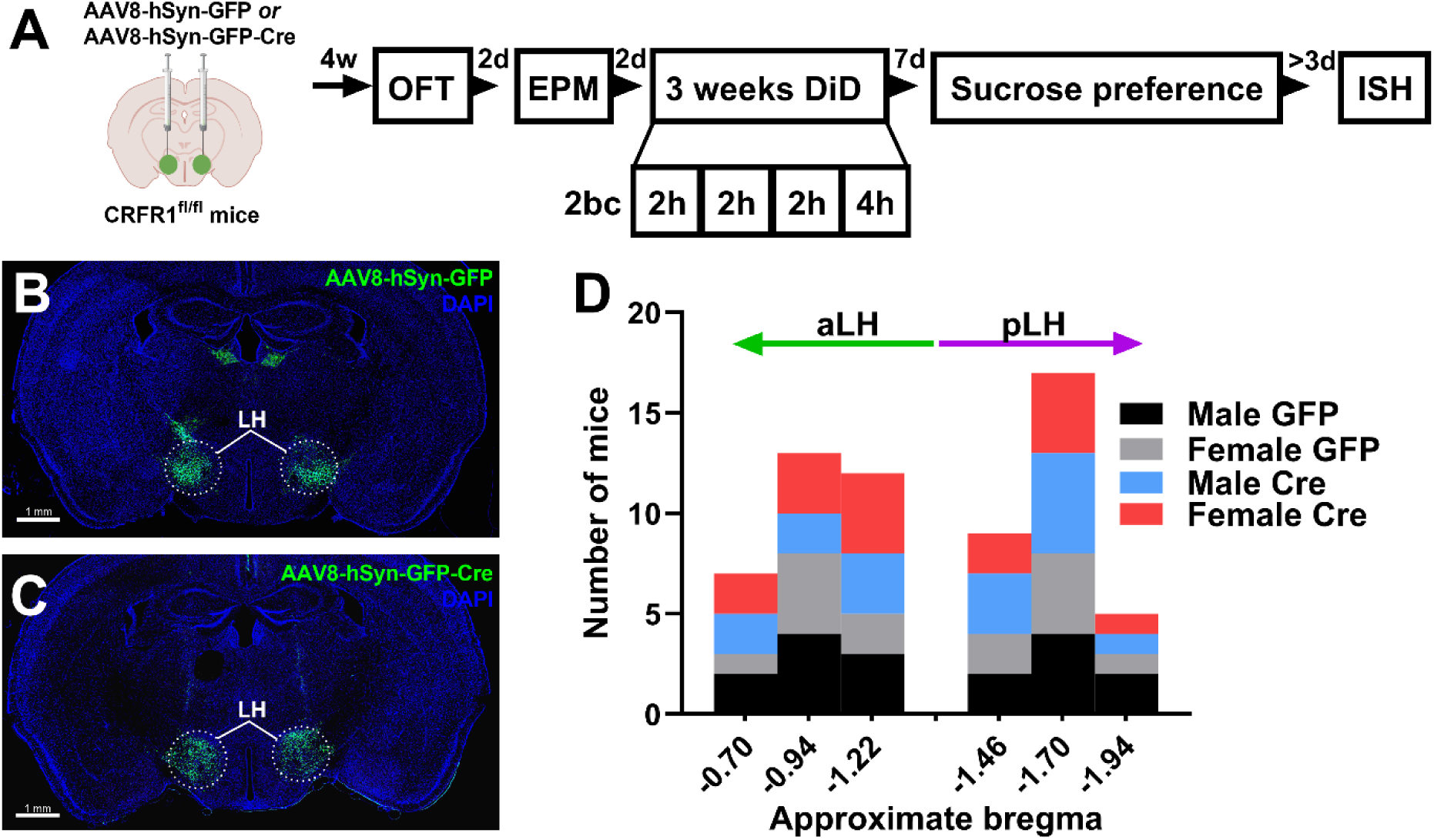
Genetically-targeted knockdown of *Crfr1* in the LH. **(A)** Experiment timeline. Male and female CRFR1^fl/fl^ mice received bilateral intra-LH infusions of GFP control virus (n = 17 male, 14 female) or Cre virus (n = 16 male, 16 female). Four weeks later, mice underwent open field test (OFT) and elevated plus maze (EPM). Mice then received 3 weeks of Drinking in the Dark (DID) consisting of 2-bottle choice (2bc) 20% alcohol drinking for 2 hours on Monday-Wednesday and 4 hours on Thursday. Sucrose preference (1% concentration) was tested 1-week following the last DID session. Brains were flash frozen for fluorescent *in situ* hybridization (FISH) >3 days after sucrose testing was complete. Representative images of **(B)** GFP- and **(C)** Cre-virus expression. Location of virus expression was cross-referenced to the Franklin and Paxinos mouse brain atlas and **(D)** mice were assigned a bregma coordinate along the anterior-posterior axis. Bregma of –1.34 was used as the center point to divide mice into anterior or posterior LH (aLH and pLH, respectively).

### CRFR1 KD in the pLH increases binge alcohol consumption

The effects of CRFR1 KD in the LH on binge alcohol consumption were assessed across 3 weeks of DID (**Fig. 2**). An omnibus ANOVA of alcohol consumption across days revealed that females consumed more alcohol than males across days, in line with previous studies^15,21,22^, and alcohol intake varied as a function of an interaction between virus and AP bregma, independent of sex (**Fig. 2A**, **2E**; 2 x 2 x 2 x 12 ANOVA; day main effect, *F*_(11,605)_ = 41.81, *p* < 0.001; sex main effect, *F*_(1,55)_ = 33.75, *p* < 0.001; AP main effect, *F*_(1,55)_ = 5.93, *p* = 0.02; day x sex interaction, *F*_(11,605)_ = 3.10, *p* < 0.001, Bonferroni post hoc female > male days 2-6, 7-12; virus x AP interaction, *F*_(1,55)_ = 11.67, *p* = 0.001). Specifically, mice expressing Cre in the pLH consumed more alcohol than their aLH GFP control or Cre counterparts (**Fig. 2A**, **2E**; Bonferroni post hoc, p < 0.05). These effects were mirrored when cumulative EtOH intake was analyzed independent of day (**Fig 2C**). Alcohol preference varied as a function of sex across days and as a function of an interaction of virus and AP bregma (**Fig 2B**, **2F**; 2 x 2 x 2 x 12 ANOVA; day main effect, *F*_(11,605)_ = 3.49, *p* < 0.001; day x sex interaction, *F*_(11,605)_ = 2.20, *p* = 0.01, male > female day 2, male < female day 9; virus x AP, *F*_(1,55)_ = 5.77, *p* = 0.02). *Post hoc* analysis revealed that, while aLH and pLH Cre groups differed independent of sex, only males showed a significant increase in alcohol preference in the pLH group compared to GFP (Bonferroni post hoc, *p* < 0.05). Females had higher BAC values following the last DID session, independent of group (Fig. 2D; ANOVA, sex main effect only, *F*_(1,43)_ = 6.54, *p* = 0.01). Correlational analysis revealed a significant negative relationship between total alcohol consumed and distance from bregma of LH Cre virus expression (**Fig. 2H**; Pearson correlation, male Cre: r(16) = −0.52, p = 0.04, female Cre: r(16) = −0.67, *p* = 0.004) that is absent for GFP controls (**Fig. 2G**; Pearson correlation, male GFP: r(17) = −0.33, *p* = 0.20; female GFP: r(14) = 0.09, *p* = 0.77). These data suggest that CRFR1 KD has divergent effects on alcohol consumption along the anterior-posterior axis, with more posterior CRFR1 KD in the LH driving increased binge drinking.

**Figure 2.**
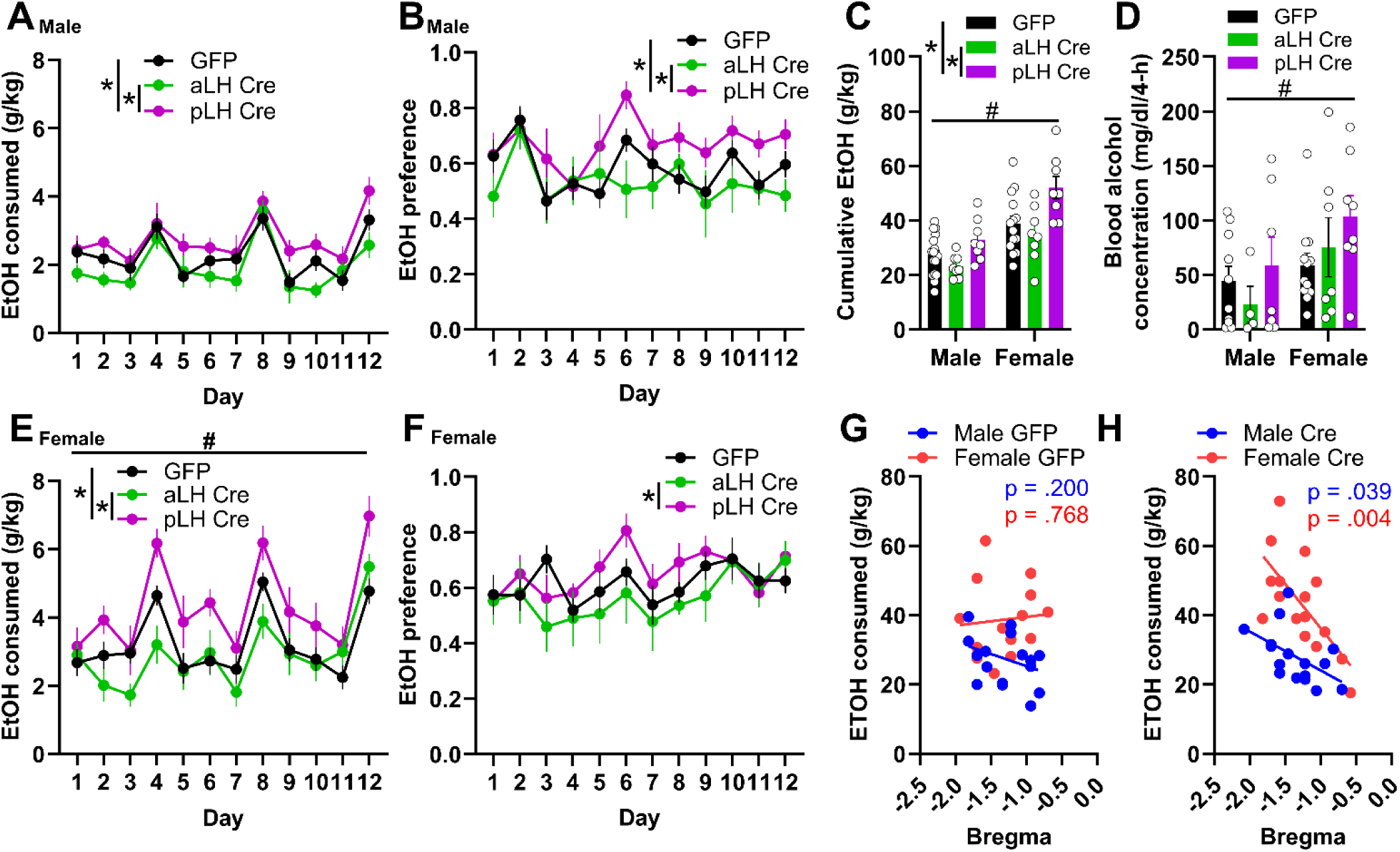
Effects of CRFR1 KD in the LH on binge alcohol consumption and preference. Male and female mice underwent three weeks of DID. Daily alcohol consumption in **(A)** male and **(E)** female mice. Daily alcohol preference in **(B)** male and **(F)** female mice. **(C)** Cumulative alcohol consumption. **(D)** Blood alcohol concentration taken after the final 4 hour DID session. Correlational analysis of total alcohol consumption and anterior/posterior (AP) distance from bregma of **(G)** GFP- and **(H)** Cre-virus expression. Data are represented as mean +/− SEM. *virus x AP interaction effect; #sex main effect; Bonferroni post hoc tests, where appropriate; All *p* < 0.05.

### CRFR1 KD blunted sucrose preference but had no effect on water consumption or body weight

Given the well-established role of the LH in feeding, drinking, and maintenance of homeostatic balance^11,12,23^, we assessed effects of CRFR1 KD on water consumption and body weight during DID and subsequently assessed sucrose preference (**Fig. 3**). In contrast to the effect of CRFR1 KD on alcohol consumption, total water consumption was higher in females independent of group (**Fig. 3A**; 2 x 2 x 2 ANOVA; sex main effect only, *F*_(1,55)_ = 11.26, *p* = 0.001). Average body weight varied by sex and as function of virus and AP bregma (**Fig. 3B**; 2 x 2 x 2 ANOVA; sex main effect, F_(1,55)_ = 132.63, *p* < 0.001; virus x AP interaction effect, *F*_(1,55)_ = 4.75, *p* = 0.03). Males were heavier than females, and body weight was not different between GFP and Cre groups. Although a marginal effect was observed when comparing aLH versus pLH Cre mice collapsed across sex, (Bonferroni post hoc tests, *p* < 0.05), Cre-mice were not different from GFP-mice (*p* > 0.05). At least 10 days after the last day of DID, mice were assayed for sucrose preference and measurements were taken at the 2- and 4-hour time points. Sucrose preference varied as a function of virus dependent on time point, independent of AP group or sex (**Fig. 3C**; 2 x 2 x 2 x 2 ANOVA; time main effect, *F*_(1,35)_ = 8.05, *p* = 0.01; virus x time interaction, *F*_(1,35)_ = 4.70, *p* = 0.04; all other *p* > 0.05). Specifically, Cre mice showed a blunted sucrose preference initially but returned to a GFP control level during the last 2 hours of the 4-hour assay (Bonferroni post hoc tests, p < 0.05). These data suggest that the divergent effects of CRFR1 KD on binge alcohol consumption based on anterior-posterior location of viral expression were not due to broad effects on neutral or rewarding fluid consumption or changes in body weight due to homeostatic imbalance.

**Figure 3.**
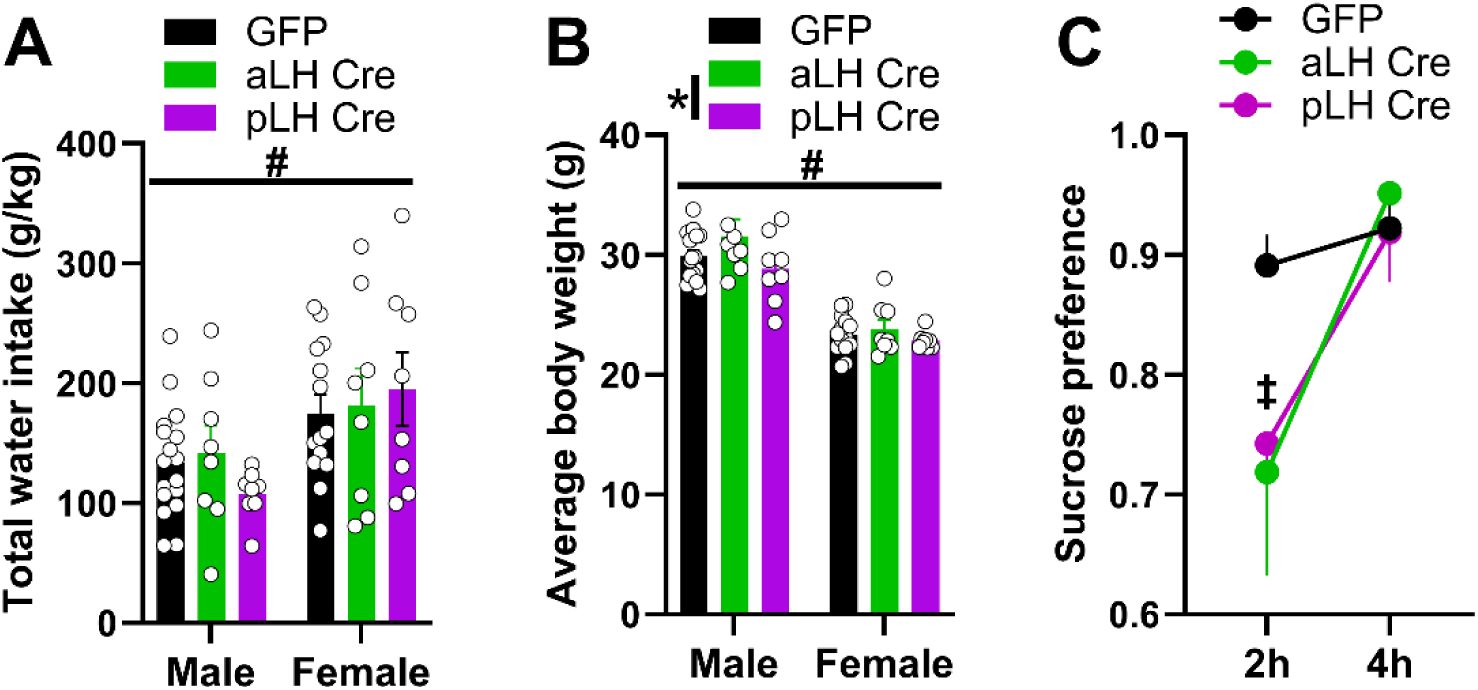
Effects of CRFR1 KD on water intake, body weight, and sucrose preference. **(A)** Total water intake and **(B)** average body weight during 3 weeks of DID sessions. **(C)** Sucrose preference measured at 2- and 4-hour time points shown collapsed across sex due to lack of sex effects. Data are represented as mean +/− SEM. #sex main effect; *virus x AP bregma interaction effect; ‡time x virus interaction effect; Bonferroni post hoc tests, where appropriate; All *p* < 0.05.

### Females exhibit higher sensitivity to the anxiogenic effects of LH CRFR1 KD in the EPM

The effects of CRFR1 KD in the LH on anxiety-like behaviors were assessed in the open-field test (OFT) and elevated plus maze test (EPM). During the OFT, center entries varied as a function of sex, virus, and AP bregma (**Fig. 4A**; 2 x 2 x 2 ANOVA, sex main effect, *F*_(1,35)_ = 5.60, *p* = 0.02; virus main effect, *F*_(1,35)_ = 8.31, *p* = 0.01; sex x virus x AP interaction, *F*_(1,35)_ = 4.16, *p* = .05). Females exhibited more center entries than males and Cre-virus mice had fewer center entries relative to GFP mice, independent of AP bregma (Bonferroni’s post hoc tests, *p* < 0.05). Center duration varied as a function of virus only, with Cre-virus mice spending less time in the center (**Fig. 4B**; 2 x 2 x 2 ANOVA; virus main effect only, *F*_(1,35)_ = 9.22, *p* = 0.004). However, correlational analysis revealed a positive correlation of center duration with location of virus expression that was only present for Cre-virus females (**Fig. 4C-D**; GFP male: r(12) = 0.265, p = 0.41, GFP female: r(9) = −0.04, *p* = 0.91, Cre male: r(11) = −0.04, *p* = 0.91; Cre female: r(11) = 0.79, *p* = 0.004). These effects were not due to a deficit in locomotion, as analysis of distance traveled revealed an *increase* in distance traveled in pLH-Cre relative to aLH-Cre mice, independent of sex (**Fig. 4E**; 2 x 2 x 2 ANOVA, virus x AP interaction only, *F*_(1,35)_ = 4.09, *p* = 0.051, Bonferroni post hoc test, aLH Cre < pLH Cre, *p* = 0.03). Time-course analysis indicated that the observed effects of CRFR1 KD in pLH-Cre mice on locomotion are due primarily to increased exploration during the first 1-minute time bin (**Fig. 4F**; 2 x 2 x 2 x 10 ANOVA, time main effect *F*_(9,315)_ = 39.17, *p* < 0.001; sex x virus x time interaction, *F*_(9,315)_ = 1.92, *p* = 0.05; virus x AP x time interaction, *F*_(9,315)_ = 2.77, *p* = 0.004; Bonferroni post hoc test, pLH bin 1 > aLH cre and GFP bin 1, *p* < 0.05). For the EPM, open arm entries were not significantly different between groups (**Fig. 4G**; 2 x 2 x 2 ANOVA, all p > 0.05). However, time spent in the open arm varied as a function of sex and virus (**Fig. 4H**; 2 x 2 x 2 ANOVA, sex x virus interaction, *F*_(1,35)_ = 4.69, *p* = 0.04). Specifically, Cre-virus females showed reduced open arm duration, independent of AP bregma, with no effect in males (Bonferroni post hoc tests, *p* < 0.05). Together, these data suggest that CRFR1 KD produced anxiety-like effects that are sex- and testing context-dependent, with females exhibiting higher sensitivity to the anxiogenic effects of LH CRFR1 KD in the EPM.

**Figure 4.**
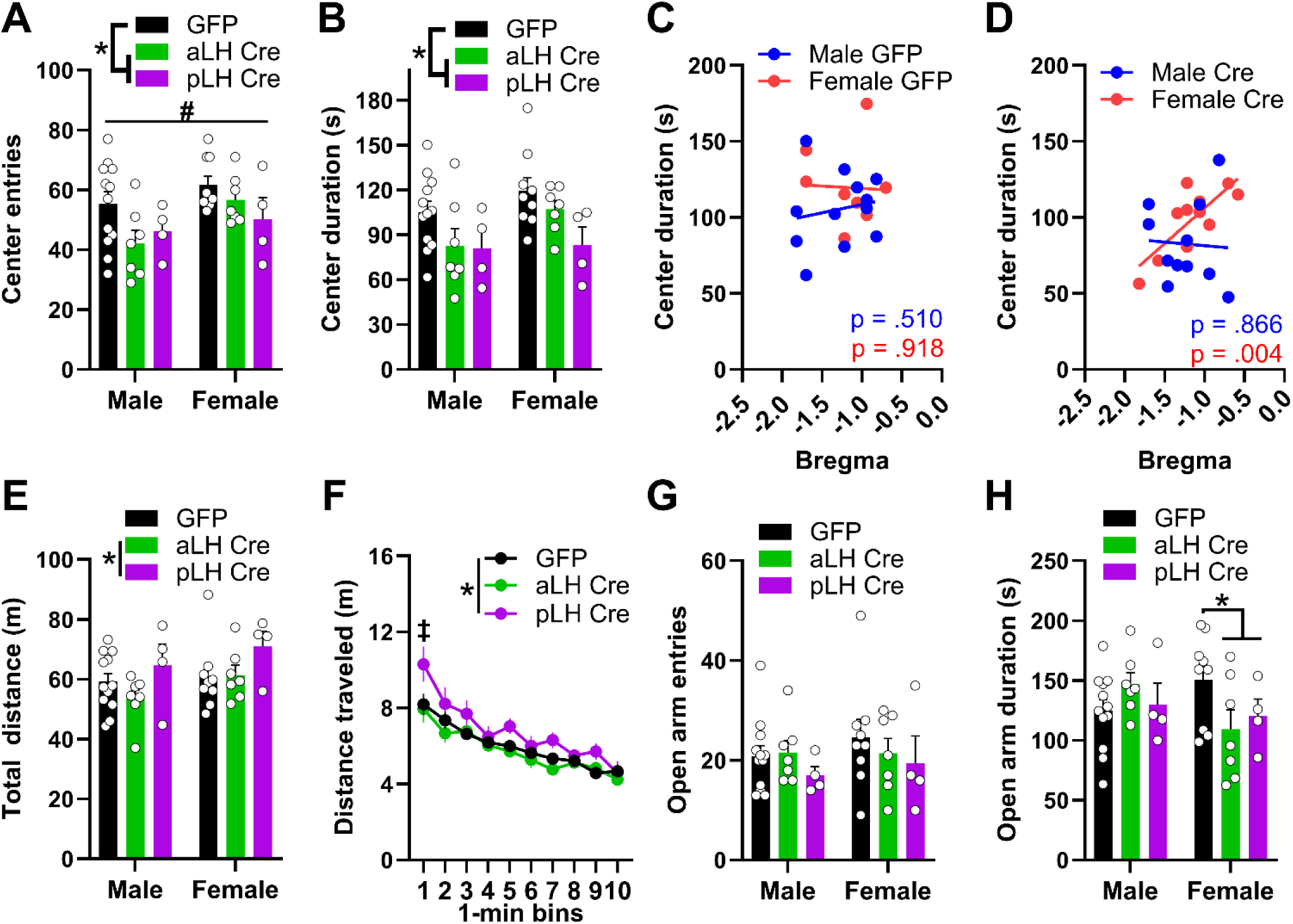
Effects of CRFR1 KD on anxiety-like and locomotor behavior. Anxiety-like behavior was assessed in the open field test (OFT) and elevated plus maze (EPM). OFT **(A)** center entries and **(B)** center duration. Correlational analysis of OFT center duration and anterior/posterior (AP) location of **(C)** GFP- or **(D)** Cre-virus expression in the LH. Distance traveled in the OFT shown as **(E)** total across session and **(F)** per 1-minute bin. EPM **(G)** open arm entries and **(H)** open arm duration. Data are represented as mean +/− SEM. *virus x AP interaction effect; #sex main effect; ‡virus x time x AP interaction effect; Bonferroni post hoc tests, where appropriate; All *p* < 0.05.

## DISCUSSION

The present study investigated the role of LH CRFR1 expression in binge alcohol consumption, anxiety-like behaviors, and sucrose preference in male and female CRFR1^fl/fl^ mice (**Fig. 1**). Knockdown of CRFR1 in the LH had divergent effects on alcohol consumption depending on the location of Cre-virus expression along the anterior-posterior axis, independent of sex (**Fig. 2**). Specifically, KD of CRFR1 in the pLH increased alcohol consumption relative to KD in aLH or GFP controls. In line with this, total alcohol consumption was negatively correlated with the AP coordinate of CRFR1 KD in the LH. These effects were not due to changes in neutral fluid consumption or energy homeostasis, as Cre mice had similar water consumption and body weight as their GFP counterparts (**Fig. 3A-B**). CRFR1 KD decreased sucrose preference independent of virus expression location in the LH (**Fig. 3C**), suggesting that the observed effects on alcohol consumption are not due to a broad shift in rewarding caloric fluid intake. Interestingly, the effects of CRFR1 KD on anxiety-like behaviors were sex- and context-specific. While male and female Cre mice showed reduced center entries and duration in the OFT, only females showed a correlation between center duration and virus placement (**Fig. 4D**). Likewise, in the EPM, only female Cre mice showed reduced open arm duration (**Fig. 4H**), suggesting that females may be more sensitive to the anxiety-like state induced by CRFR1 KD in the LH. These data are the first to report on the effects of chronic CRFR1 disruption in the LH on binge alcohol consumption and anxiety-like behaviors. Our novel finding that the effects of CRFR1 KD diverge along the anterior-posterior axis in the LH provides an impetus for future investigations into potential cell-type or circuit-specific function of the LH in regulating binge drinking.

There is considerable but inconsistent literature characterizing the effects of CRFR1 manipulations on binge drinking. Centrally administered CRF blunts alcohol intake^24^; yet, surprisingly, there is also a reduction in alcohol intake following central CRFR1 antagonism^9^. These conflicting data could suggest separate neural substrates are mediating the two effects. Alternatively, CRF likely also elicits effects through CRF receptor 2 (CRFR2). In the VTA, intact CRFR2 signaling is necessary for the effects of CRFR1 antagonism on binge alcohol consumption^25^. We previously showed that acute CRFR1 antagonism in the LH decreased binge drinking in males while CRFR2 agonism had no effect^15^, suggesting CRFR1 may be the dominant CRF receptor in LH. In contrast, CRFR1 and CRFR2 antagonism in the LH similarly normalizes stress-induced anxiety-like behavior, indicating a pro-stress role for both receptors^26^. It may be that chronic knockdown of CRFR1 produced a compensatory increase in CRFR2 signaling, or perhaps other modulatory systems in the LH.

Both CRFR1 and CRFR2 are expressed in the LH at densities that vary along the anterior-posterior axis, with CRFR2 and CRFR1 expressed more densely in the aLH and pLH, respectively^27^. Interaction between these receptor systems at different levels throughout the LH may play a role in binge alcohol consumption. However, prior pharmacological experiments have not differentiated between the aLH and pLH^15^. In the present study, we used genetically targeted KD of CRFR1 in the LH using CRFR1^fl/fl^ mice. An important follow-up investigation would be to repeat this experiment using CRFR2-flox or CRFR1/CRFR2 double-flox mice to provide further insight into the role of these receptors in binge alcohol drinking.

The effect of chronic CRFR1 KD in the LH on binge alcohol consumption was independent of sex, in contrast to prior reports using pharmacology and DREADD approaches. We recently reported that acute treatment with a CRFR1 antagonist in the LH blunts binge alcohol consumption selectively in males^15^. Similarly, chemogenetic disruption of a CeA CRF to LH circuit decreased alcohol consumption in males only. One possibility is that CRFR1 antagonism in the LH acutely alters aspects of alcohol consumption distinct from those altered by chronic knockdown and that preferentially affected males. Female mice show increased motivation for alcohol, responding more on an operant lever for alcohol than males^28^. Females also show greater resistance to quinine adulteration than males, which coincides with higher c-fos activation in the VTA^21^. Together, these data suggest that females may be more sensitive to the rewarding properties of alcohol than males, rendering them resistant to disruption by an acute pharmacological manipulation. In contrast, chronic disruption of CRFR1 signaling in the LH may overcome this sex difference to produce sex-independent effects on binge alcohol consumption. Alternatively, the chronic deletion of CRFR1 may lead to compensatory changes in other receptors within the LH or downstream brain regions. Our prior work assessed a single dose of either the CRFR1 antagonist or DREADD agonist. It remains possible that higher doses of antagonist could be effective to reduce drinking in females.

Only females displayed an anxiety-like response in the EPM to CRFR1 KD in the LH. Additionally, correlational analysis of OFT center time revealed that only females showed increasing anxiety-like behavior with more posterior Cre-virus expression in the LH. These data provide novel evidence that the pLH may be a site of sexual dimorphism for the regulation of anxiety-like states. To date, there is no literature characterizing sex differences in the role of the LH in anxiety. Our OFT data contrast with anxiolytic effects of CRFR1 antagonism in the LH of male rats^29^, possibly due to antagonist action on pre-synaptic CRFR1. Interestingly, LH output through activation of vGat or galanin expressing LH output neurons is anxiolytic^23,30^. Therefore, our anxiogenic effect of CRFR1 KD could be due to decreased activation of these neuronal populations in the LH. Our sex-dependent results are interesting in light of the human literature indicating that women exhibit a higher prevalence of anxiety disorders and increased activation of the LH following a stressor ^31,32^. The present data suggest anatomical differences in LH CRFR1 regulation of anxiety-like behavior and hint at a neurophysiological mechanism underlying sex differences in stress and anxiety processing.

The effects of CRFR1 KD on binge alcohol consumption diverged based on the location of Cre-virus expression in the LH along the anterior-posterior axis. Though differences in ascending projections of the aLH and pLH are long established^16^, only recently have rigorous investigations into functional differences in LH substructures been reported. The two populations of neurons within the LH that have received the most attention for their roles in motivated behavior are inhibitory vGat and excitatory vGlut2 neurons, which have distinct output targets and functions^33,34^. vGlut2 neurons are of particular relevance to our findings because they exhibit a diverging pattern of expression and output targets along the anterior-posterior axis, with aLH and pLH vGlut2 neurons primarily projecting to the LHb and VTA, respectively^17^. While both LH vGlut2 circuits encode rewarding and aversive tastants^17^, they are distinct in how they encode reward signals based on satiety (i.e., motivational) state. Our CRFR1 KD in the LH may have had divergent effects along the anterior-posterior axis by altering distinct output targets of the LH, shifting the motivational salience of alcohol reward. However, a limitation of our genetically-targeted KD is a lack of circuit^35^ specificity, making interpretation challenging in this regard. It would be interesting to investigate the role of distinct populations of CRFR1-expressing LH neurons using a more targeted approach. Though not assessed here, the LH also varies considerably in connectivity and gene expression along the dorsal-ventral axis^36,37^, which may add additional nuance to interpretation as the role of the LH in binge drinking is assessed moving forward.

### Conclusions

Historically, the LH has primarily been characterized as a feeding center. There is a limited literature highlighting LH functional and neurocircuit diversity along the anterior-posterior axis^17,33^. The present study builds on this work, providing the first evidence that CRFR1 regulates binge alcohol intake in a manner dependent on AP location in the LH. These effects were not due to changes in homeostatic pressures, and were not mirrored in sucrose preference, suggesting a novel, alcohol-specific role of LH CRFR1 in binge alcohol consumption. Notably, the LH regulates the rewarding properties of other drugs of abuse, including cocaine^38^ and morphine^35^, but the role of CRFR1 has not been assessed in these contexts. Recent evidence has prompted a reconceptualization of LH function outside of its role in energy homeostasis, instead attributing to the LH a broad role as a central hub for the regulation of motivated behavior and processing of reward-proximal information^39^. This raises important questions considering the present data as to what facet of alcohol intake is regulated by CRFR1 signaling in the LH, and how this might change with chronic alcohol misuse. Future investigations into alcohol-specific function of LH neurocircuitry and CRFR1 signaling will be critical to understand how these factors contribute to the development or progression of AUD.

## Declaration of Interests

The authors have nothing to declare.

## Funding Sources

This research was supported by grants from the National Institutes of Health’s (NIH) R01AA022048 and T32AA007573.

---

**Supplemental Figure 1.**
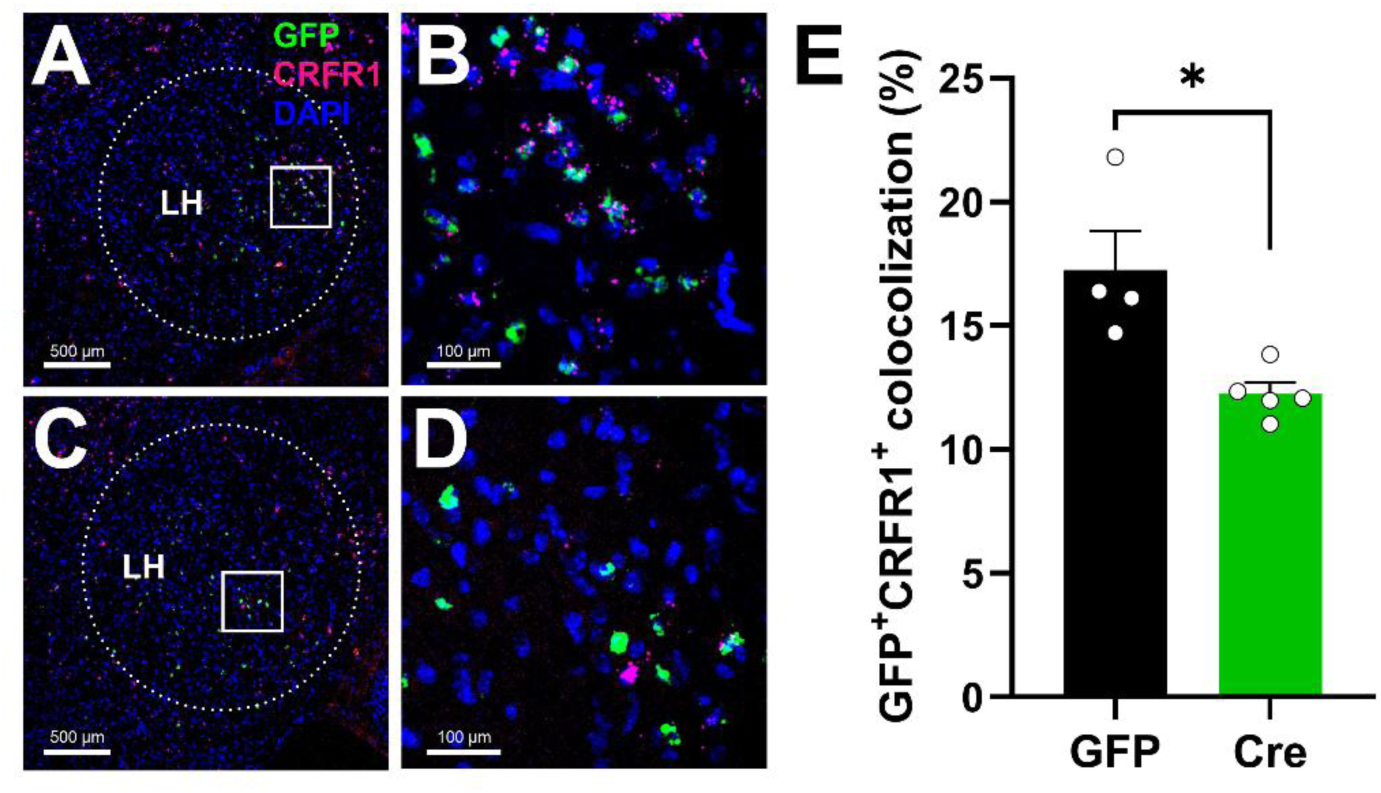
Efficacy of viral knockdown was assessed using FISH labeling *GFP* and *Crhr1* mRNA in tissue sections containing the LH of **(A)** GFP- and **(C)** Cre-virus expressing mice. **(B**, **D)** 20x magnification of the LH (area in white square). **(E)** Quantification of *GFP*^+^*Crhr1^+^* colocalized cells revealed a ∼29% reduction in colocalized cells of Cre mice relative to GFP mice (n = 16 slices from 9 mice). Data are represented as mean +/− SEM. *t-test, different from GFP, *p* = 0.04.

## Notes

### Competing Interest Statement

The authors have declared no competing interest.

